# Type 2 Diabetes: One Disease, Many Pathways

**DOI:** 10.1101/648816

**Authors:** Joon Ha, Arthur Sherman

**Affiliations:** Laboratory of Biological Modeling, National Institutes of Health, Bethesda, MD USA

## Abstract

Diabetes is a chronic, progressive disease that calls for longitudinal data and analysis. We introduce a longitudinal mathematical model that is capable of representing the metabolic state of an individual at any point in time during their progression from normal glucose tolerance to type 2 diabetes (T2D) over a period of years. As an application of the model, we account for the diversity of pathways typically followed, focusing on two extreme alternatives, one that goes through impaired fasting glucose (IFG) first, and one that goes through impaired glucose tolerance (IGT) first. These two pathways are widely recognized to stem from distinct metabolic abnormalities in hepatic glucose production and peripheral glucose uptake, respectively. We confirm this but go beyond to show that IFG and IGT lie on a continuum ranging from high hepatic insulin resistance and low peripheral insulin resistance to low hepatic resistance and high peripheral resistance. We show that IFG generally incurs IGT, and IGT generally incurs IFG on the way to T2D, highlighting the difference between innate and acquired defects and the need to assess patients early to determine their underlying primary impairment and appropriately target therapy. We also consider other mechanisms, showing that IFG can result from impaired insulin secretion, that non-insulin dependent glucose uptake can also mediate or interact with these pathways, and that impaired incretin signaling can accelerate T2D progression. We consider whether hyperinsulinemia can cause insulin resistance in addition to being a response to it and suggest that this is a minor effect.

## Introduction

Diabetes is by definition a state of hyperglycemia, but its natural history is diverse. For example, some individuals experience fasting hyperglycemia first (Impaired Fasting Glucose, IFG), followed by hyperglycemia at the two-hour time point (2hPG) of an oral glucose tolerance test (OGTT), defined as Impaired Glucose Tolerance (IGT), and some experience these in the opposite order. Eventually, all people with diabetes will have both IFG and IGT (combined glucose impairment, CGI), so we refer to these two pathways as “IGT-first” and “IFG-first”, respectively. An important implication of these observations is that the best period for determining differences in the underlying physiology of these pathways is during the pre-diabetic stage, when the phenotypes are still distinct.

Prediabetes is also the stage in which progression to type 2 diabetes (T2D) can be markedly delayed or prevented (45), and interventions can plausibly be made even more effective by targeting the specific metabolic defects of the patient. For example, IFG is generally thought to reflect insulin resistance at the liver, resulting in elevated hepatic glucose production (HGP), whereas IGT is thought to reflect peripheral insulin resistance, mainly in muscle, resulting in reduced glucose disposal. One would like to know whether using drugs that primarily affect hepatic or peripheral insulin resistance makes a material difference for patients on the IFG-first or IGT-first pathways, and whether any such benefit carries over once T2D has begun.

Moreover, diabetes is not a state that one enters and exits, like an infection, but a chronic condition that is the culmination of a series of gradual changes. Understanding of this progression is best obtained by longitudinal studies over a period of years or decades. Here we will focus on one such study, the Baltimore Longitudinal Study of Aging (BLSA) (51), which asked whether CGI is an obligatory stage between IFG and T2D and between IGT and T2D (Supplemental Fig. S1; all supplemental material is available at Figshare, https://doi.org/10.6084/m9.figshare.10792412).

These features of diabetes suggest that a longitudinal mathematical model for disease progression could be a valuable adjunct to clinical studies. Here we establish such a model and demonstrate that it can distinguish hepatic and peripheral insulin resistance and can simulate the IGT-first and IFG-first pathways to T2D over a period of years. We will use the model to address the clinical questions raised by the BLSA and Perrault studies. By tracking virtual patients continuously in time, the model can interpolate between clinical observations, which are necessarily sparsely sampled, and indicate what is likely to have happened in the interim.

The model provides broader insight into the different insulin-resistance phenotypes, showing that a wide array of pathways, ranging from isolated IFG to isolated IGT to combinations of the two, can be obtained by combining different degrees of hepatic and peripheral insulin resistance. Thus, the apparent clinical diversity of individual paths comprises a set of quantitative variants within a unified process of metabolic dysfunction.

There is also diversity in the relative contributions of insulin resistance and beta-cell dysfunction to diabetes pathogenesis. This is particularly apparent in comparing diabetes risk factors in populations with a high prevalence of obesity, such as those of African descent and Native Americans, to populations in which diabetes risk is seen among lean individuals and beta-cell function is weaker, such as South and East Asians; populations of European descent tend to lie in between (46). Furthermore, beta-cell dysfunction can be subdivided into defects in first- and second-phase insulin secretion. As a first installment on this large and complex set of problems, we examine whether the loss of first-phase secretion early in pre-diabetes plays a fundamental causal role in T2D development or is just a useful marker.

### General modeling approach

The diabetes field is fortunate to have a strong tradition of mathematical modeling (20, 49). A common feature of those models is that they provide snapshots in time of the metabolic state of an individual, including insulin resistance and beta-cell function. These models have been used to track progression by taking a series of snapshots, but they do not contain mechanisms of progression and do not describe trajectories of progression.

The model developed here belongs to a different family of models that seek to explain disease progression mechanistically, rather than assess the current state. It is an extension of our previously published model of beta-cell mass and function, which successfully simulated progression to diabetes in rodents over months and humans over years (37). That model, in turn, was based on the seminal model of (75), which expressed mathematically the hypothesis that beta-cell mass provides negative feedback on a slow time scale to compensate for insulin resistance. If that compensation is inadequate, however, the toxic effects of very high glucose overcome the stimulatory effects of moderately elevated glucose. The normal homeostatic negative feedback is converted to positive feedback, leading to deterioration in glucose tolerance and culminating in diabetes. This fundamental concept has been incorporated into other models that broadly agree but emphasize different details (23, 24, 35, 38, 43, 79). Notably, the model of (38) was shown to be able to account for the progression of fasting hyperglycemia and T2D observed in the Diabetes Prevention Program (DPP)(45).

A third class of modeling studies has fit longitudinal data from clinical studies with non-linear mixed effects statistical approaches to assess the magnitude of treatment effects on glucose, insulin and HbA1c (18, 25) or, by fitting to a modified form of the model of (75), on beta-cell mass and insulin sensitivity (67).

See (24) for further comment on the models cited here and other models, and see (54) for a perspective on modeling T2D.

The initial wave of mechanistic longitudinal models simulated fasting or average daily glucose and were therefore unable to describe post-prandial responses or responses to glucose challenges such as OGTTs and IVGTTs. A recent model (24) added the capability of following daily and post-challenge glucose variations and was fit to OGTT data from the DPP. It also introduced a distinction between peripheral and hepatic insulin resistance in order to account for the effects of drug and lifestyle interventions on these parameters.

We also, using a different methodology, introduce simulations of OGTTs at selected points during progression, and separate representations for hepatic and peripheral insulin resistance. We use these new features to differentiate progression by either 2hPG or FPG. We perform IVGTTs as well to illustrate the evolution of the acute insulin response to glucose (AIRg), often used to assess beta-cell function. This requires the model to simulate first and second phase secretion, which we accomplish by incorporating a previously published model for insulin granule exocytosis (17).

This study focuses on mechanism and insight, rather than assessment. Instead of fitting parameters to particular data sets, we assume parameters and investigate the trajectories of glycemia and insulin secretion that result. We demonstrate that the model captures known features of diabetes pathogenesis data and provides novel insights and interpretations of the data. We conceptualize this as building a factory, not a product. Once the utility of the model is established, we expect that a wide variety of applications to clinical data will become possible.

## Materials and Methods

We briefly review the previous version of the model (37) and then describe the enhancements introduced here. The enhanced version has been used to study clinical implications of differences in glucose time courses during an OGTT (19), but was not documented in detail. Parameter values and details of functions not given here are in the Appendix (Supplementary Material).

### Previous version of the model

The core element, retained in the new version, is a glucose-insulin feedback loop, represented by two differential equations adapted from the Minimal Model of Bergman and Cobelli (8), who built on models going back to the 1960’s (2, 11), as modified in (75):

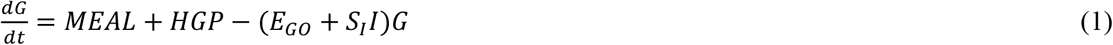

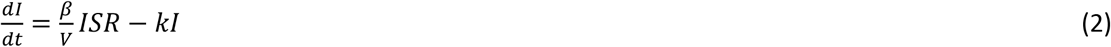

The glucose (*G*) equation (Eq. 1) says that *G* increases on a time scale of hours as a result of meal influx and hepatic glucose production (*HGP*) and decreases as a result of uptake, which has both insulin-independent and insulin-dependent components. The factor *S*_*I*_ in the insulin-dependent term is closely related to the well-known sensitivity to insulin reported by the Minimal Model. The insulin (*I*) equation (Eq. 2) says that *I* decreases due to removal, mainly in the liver, with rate constant *k*, and increases due to secretion by beta cells, where *β* is the beta-cell mass, *ISR* is the insulin secretion rate per unit mass, and *V* is the volume of distribution. In the original version, *ISR* depended only on *G*, through the rate of beta-cell metabolism *M*:

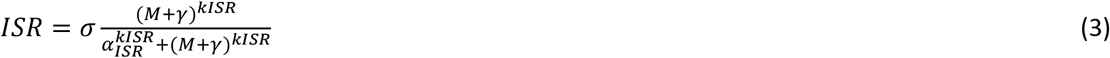

where *M* was assumed to be a sigmoidally-increasing function of *G*:

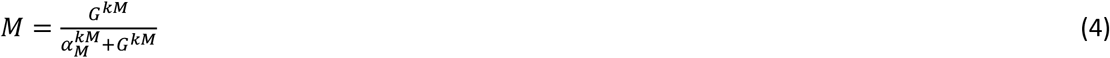

The parameter *γ* in Eq. (3) represents the effect of K(ATP) channel density to shift the glucose dependence of secretion (the triggering pathway (39)); when the channel density is low, *γ* is high, and shifts the dependence to the left, increasing secretion for the same level of *M* because Ca^2+^ is higher. Experiments in (34) showed that mouse beta cells in vitro adjust the K(ATP) channel density down in response to sustained (overnight) elevated glucose. This has also been observed in vivo in humans (41, 42, 44), along with evidence of reduced insulin clearance, *k*, (19–21). For the most part, we omit this for simplicity, but we consider the implications of reduced clearance towards the end of Results. This can be viewed as the first line of defense through enhanced beta-cell function against insulin resistance over a time scale of days (e.g. holiday overeating).

The value of *γ* depends on glucose, which is taken into account by adding a third differential equation to the system:

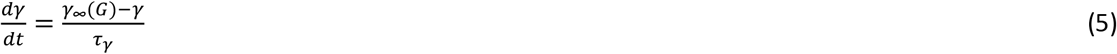

where *γ*_∞_ is an increasing sigmoidal function of *G*, and *τ*_*γ*_ is the time constant.

Insulin resistance that persists over longer periods (months in humans) despite reduced K(ATP) channel density, is assumed to trigger a further level of compensatory increased beta-cell function via σ, the maximal insulin secretion capacity (Eq. 3). This corresponds to the amplifying effects of metabolism and/or modulators such as GLP-1 and ACh on the efficacy of Ca^2+^ to drive insulin granule exocytosis. This second aspect of beta-cell functional compensation entails a fourth differential equation:

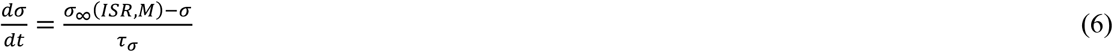

We assume that increased *ISR* (workload in the sense of (64)) leads to an increase in σ whereas increased *M* leads to a decrease in σ.

The slowest and final form of compensation for insulin resistance is increased beta-cell mass, *β*, which develops over years in humans. We assume that *β* is increased by proliferation, *P*, and decreased by apoptosis, *A*. Following the data of (64), we assume that *P* increases when *ISR* increases. We further assume that apoptosis is largely driven by metabolic stress (e.g. through increased production of reactive oxygen species) when glucose is high, so we make *A* an increasing function of *M*:

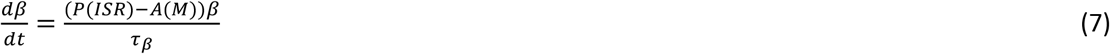

The parameters defining *P* and *A* are chosen such that modest increases in *G* result in a net increase in *β*, but large increases in *G* result in a net decrease in *β*. As in the predecessor model (75), this leads to a shift from compensation (negative feedback) to decompensation (positive feedback). In (37), we showed that this can account for the threshold behavior observed in both rodents and humans, that is, nearly steady *G* followed by a sharp, essentially irreversible increase (50, 73). We proposed this as an explanation for why prevention of T2D is much easier than reversing it once it is established. The same dynamic properties carry over in this study with the model enhanced as described next.

### New features in the model

#### Modeling glucose flux during daily meals and glucose tolerance tests

In the previous version of the model (37), *G* represented average daily glucose and insulin levels in response to steady glucose input. To address IFG and IGT, we need to be able to dissociate fasting glucose from post-challenge glucose. The first step is to introduce variable glucose influx from meals, represented by the term *MEAL* in Eq. (1). Timing of meals is standardized to 6:00 AM, 12:00 Noon, and 6:00 PM. The expression we use is:

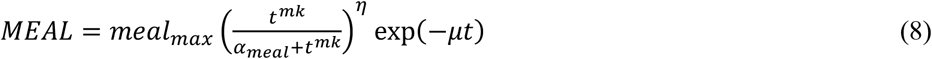

and Fig. S3 shows the glucose flux (panel A) and the resulting plasma glucose concentrations (panel B). We do not yet account for other nutrients (amino acids, fats). Flux during an OGTT is modeled with a more rapid rise and decay of flux (Eq. (A2) and shown in Fig. S3, C, D). Flux during an IVGTT rises and decays still more rapidly.

#### Modeling hepatic glucose production (HGP)

In order to distinguish peripheral and hepatic insulin resistance and describe how they are related to each other, we need to refine the model description of hepatic glucose production (*HGP* in Eq. 1). In the first version of the model (37), *HGP* was assumed to be constant, which is an acceptable approximation as long as fasting plasma insulin is adequate to compensate completely for any hepatic insulin resistance. To study the failure of compensation, however, we need to make *HGP* a decreasing function of *I*:

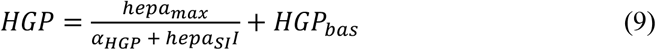

This is sufficient to give the typical drop and recovery of *HGP* after a meal (Fig. S4A).

Less obvious, but also important, we need to account for the correlation between hepatic and peripheral insulin resistance, which we do by making two of the parameters in Eq. 8, *hepa*_*max*_ and *α*_*HGP*_, functions of *S*_*I*_:

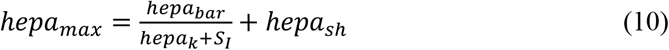

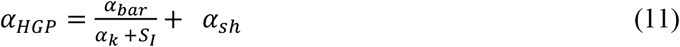

Both *hepa*_*max*_ and *α*_*HGP*_ decrease with *S*_*I*_, as shown in Figs. S4B, C, and the net effect is that *HGP* decreases as *S*_*I*_ increases (Fig. S4D).

If this, or something like this, is not done, then severe peripheral resistance combined with strong compensatory insulin secretion can result in fasting hypoglycemia, which is not the typical pattern (see Fig. S5).

In more typical cases of progression to pre-diabetes or diabetes, the relative impairment in insulin secretion would mask this effect: the level of glycemia would be reduced but hypoglycemia would not result. We chose the parameters such that HGP remains normal when insulin is elevated unless there is a defect in beta-cell mass or function.

Figure S6, E – H shows the response of HGP and glucose disposal to a simulated hyperinsulinemic, euglycemic clamp, with steady-state values in good agreement with experimental data (6).

To represent hepatic insulin resistance over and above the component related to peripheral insulin resistance, we decrease the parameter *hepa*_*SI*_ in Eq. A4, which increases HGP at any value of *I*.

Equations 9 – 11 are phenomenological expressions that achieve the goals of making *HGP* decrease with *I* and increase when insulin resistance is present, but other expressions could also possibly work. We considered making the value of *hepa*_*SI*_ depend on *I* as an alternative way to avoid hypoglycemia, but this did not work well. It is also possible that glucagon, which is not in the model, would help to prevent hypoglycemia (see *Limitations of the study* in the Discussion).

#### Modeling insulin granule exocytosis

To study the dynamics of glucose and insulin under glucose challenges such as meals, OGTT, and IVGTT, the model needs to account for the multiple kinetic components of insulin secretion. We adapted an existing model of insulin granule exocytosis (17), which was designed to capture the biphasic pattern of *ISR* in response to a glucose step in vitro or a hyperglycemic clamp in vivo. The first phase is characterized by a sharp peak of *ISR* during the first 10 minutes and the second phase by a steady increase of *ISR* over the next hour. Figure S6 shows a simulated OGTT (panels A, B) and a simulated IVGTT (panels, C, D) compared to experimental data.

These phases are mediated by progression of vesicles through a sequence of stages culminating in exocytosis (fusion with the plasma membrane and release of insulin to the circulation; Fig. S2A). A large reserve pool, treated as an inexhaustible reservoir, feeds the docked pool (vesicles at binding sites on the plasma membrane). Once docked, vesicles are primed and enter the readily releasable pool (RRP). Primed vesicles join the immediately releasable pool by becoming closely associated with voltage-dependent Ca^2+^ channels. First-phase secretion depends mainly on the size of the RRP at basal glucose, and second phase secretion is controlled by the rate of mobilization of vesicles to the docked pool.

The insulin secretion rate *ISR* in the first version (37) of the model (Eq. 3) is in the new version no longer a function of glucose, but is calculated as an output of the exocytosis model (Eqs. A11), *ISR* thus now depends on the history of exposure to *G*, as it should, not just the current value. The exocytosis model requires as input the cytosolic Ca^2+^ concentration, which is modeled as a sigmoidal function of the beta-cell metabolic rate *M* (Eq. A7), and the much higher Ca^2+^ concentration in the microdomains of Ca^2+^ channels, which is modeled as a function of cytosolic Ca^2+^ (Eq. A8). The dose response curve shift *γ*, previously included in Eq. 3, represents the dynamic changes in K(ATP) channel density as before, but now explicitly alters cytosolic Ca^2+^ at a given level of *M* (Eq. A7). Cytosolic calcium enhances the rates of mobilization of the reserve pool to the plasma membrane (Eq. A10) and priming of docked vesicles (*r*_2_ in Eqs. A12), whereas vesicle fusion is primarily controlled by microdomain Ca^2+^. The amplifying effect of glucose (39) is incorporated as a multiplicative factor in the rate of vesicle mobilization (*G*_*F*_ in Eq. A9). The effect of the incretin GLP-1and drugs that mimic it to increase insulin secretion is modeled as enhancing the amplifying effect of glucose by increasing the parameter *G*_*Fmax*_ in Eq. A9 and as enhancing vesicle priming by increasing the parameter 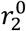 in Eq. A12. This is in accord with a key feature of these agonists, that they have little or no effect on secretion in the absence of glucose, minimizing the risk of hypoglycemia that plagues sulfonylureas. Fig. S2C shows the effect of a two-fold increase in the incretin effect on insulin secretion rate during a hyperglycemic clamp, in agreement with the effect of the GLP-1 receptor agonist liraglutide in the RISE study (68). Modeling the effect of incretins on glucose is more difficult, as it would require including their effects to inhibit glucagon secretion or promote weight loss, which is not feasible in the current model.

For IVGTT and hyperglycemic clamp simulations, we reduce *G*_*F*_ by about a factor of two to represent the lack of the incretin effect on vesicle mobilization and reduce 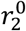 (Eq. A12) about 10-fold to represent the lack of incretin effect on vesicle priming. The rate of mobilization (Eq. A11) is also assumed to be proportional to the variable σ (Eq. 6), which thus controls the magnitude of second-phase insulin secretion. As a consequence, *ISR* implicitly includes σ as a multiplicative factor, as in Eq. (3) of the simpler, first version of the model (37), and the dynamic evolution of the maximal secretory capacity over long time scales (months) is essentially equivalent.

#### Equations for insulin sensitivity

With one exception we model peripheral insulin sensitivity as an intrinsic property of the target tissues, muscle, liver and adipose, determined implicitly by obesity and genetics. In the model, this makes it an external, time-dependent parameter of the system, and we examine how hyperinsulinemia compensates for insulin resistance. We model insulin sensitivity, *S*_*I*_, as an exponentially decreasing function of time, following (75):

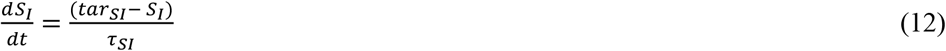

In Fig. 9, we consider a hypothesis under current discussion in the literature (35, 65, 71, 74), that insulin resistance may be caused by hyperinsulinemia, rather than the other way around. We think it is implausible that this is the sole cause of insulin resistance, and propose instead that the core component of insulin resistance is due to obesity and genetics but modified by the level of insulin *I* as a result of the negative feedback generated in the insulin signaling cascade (70). We represent the effect, which we term *induced resistance*, as follows:

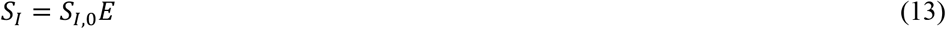

where

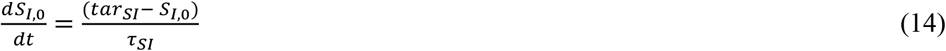

and

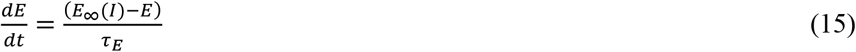

Equation (15) says that *E* relaxes towards *E*_∞_with a time constant *τ*_E_, where *E*_∞_ is a sigmoidally decreasing function of *I*,

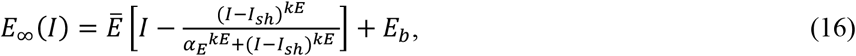

several examples of which are shown in Fig. 9F. The time constant *τ*_*E*_ is set to two days so that *E*, and hence *S*_*I*_, will respond to sustained hyperinsulinemia, not to normal insulin fluctuations following meals or glucose tolerance tests. The effect is propagated into hepatic insulin resistance through the terms containing *S*_*I*_. Other parameters are in the legend for Fig. 9 and Table S17.

### Criteria of pre-diabetes and diabetes

Following ADA criteria, we define IFG as FPG > 100 mg/dl but < 126 mg/dl, IGT as 2hPG > 140 mg/dl but < 200 mg/dl, and T2D as FPG ≥ 126 mg/dl or 2hPG ≥ 200 mg/dl. Combined glucose impairment (CGI) is defined as co-occurrence of IFG and IGT.

### Software

The model equations are solved using xppaut (5) and Matlab (The Mathworks, Natick, MA). Input files defining the parameters and initial conditions are available on Figshare at: https://doi.org/10.6084/m9.figshare.10792412).

## Results

In Figs. 1 – 4, we carry out longitudinal simulations over a period of five years starting in the NGT state in which either peripheral insulin resistance is dominant, in which case the first stage of hyperglycemia is IGT, or hepatic insulin resistance is dominant, in which case the first stage of hyperglycemia is IFG. These assumptions are applied by modeling peripheral and hepatic insulin sensitivity as exponentially decreasing, using Eqs. A13 and A14, respectively. The initial values and rates of decline are the same for all figures, but the steady state (target) values, *tar*_*SI*_ and *tar*_*hepaSI*_, are varied. The simulations calculate the daily responses to meals, but we plot only the results of OGTTs carried out periodically over the five-year time span by pausing the longitudinal simulation. The corresponding peak postprandial glucose is shown in Fig. S7. The capacity of beta-cell function to compensate for insulin resistance is assumed to be limited, except in Fig. 2. The defect consists of a right shift in *γ*_∞_ relative to Fig. 2, which can be interpreted as a slight gain of function mutation in KCNJ11, the Kir6.2 component of the KATP channels. Alterations in *σ*_∞_ representing a mild defect in insulin granule mobilization would have a similar effect.

**Figure 1:**
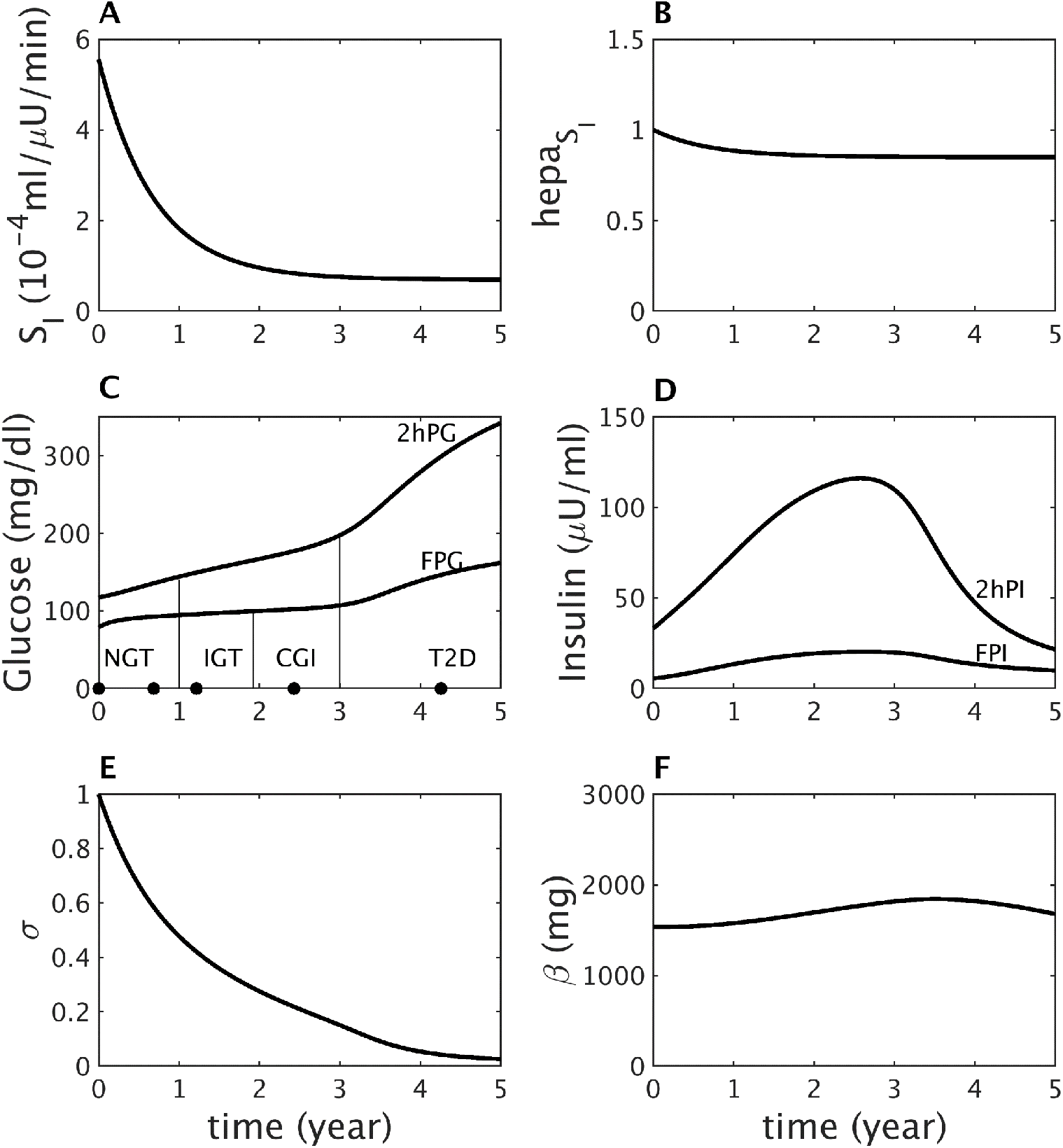
IGT-first pathway to diabetes. (A) Assumed severe decline in peripheral insulin sensitivity. (B) Assumed mild decline in hepatic insulin sensitivity. (C) Simulated longitudinal changes based on the assumptions in (A) and (B) in fasting plasma glucose (FPG) and two-hour glucose (2hPG) during OGTTs performed at each time point. The virtual subject experiences first high two-hour glucose (IGT), then high fasting glucose (CGI), and finally crosses the 2hPG threshold for T2D. (D) Simulated longitudinal changes in fasting plasma insulin (FPI) and two-hour insulin (2hPI) during the OGTTs. Insulin increases early on but decreases later. (E) The component of *β*-cell function represented by σ decreases progressively throughout. (F) The *β*-cell mass, *β* first increases during prediabetes, then decreases after diabetes onset.

**Figure 2:**
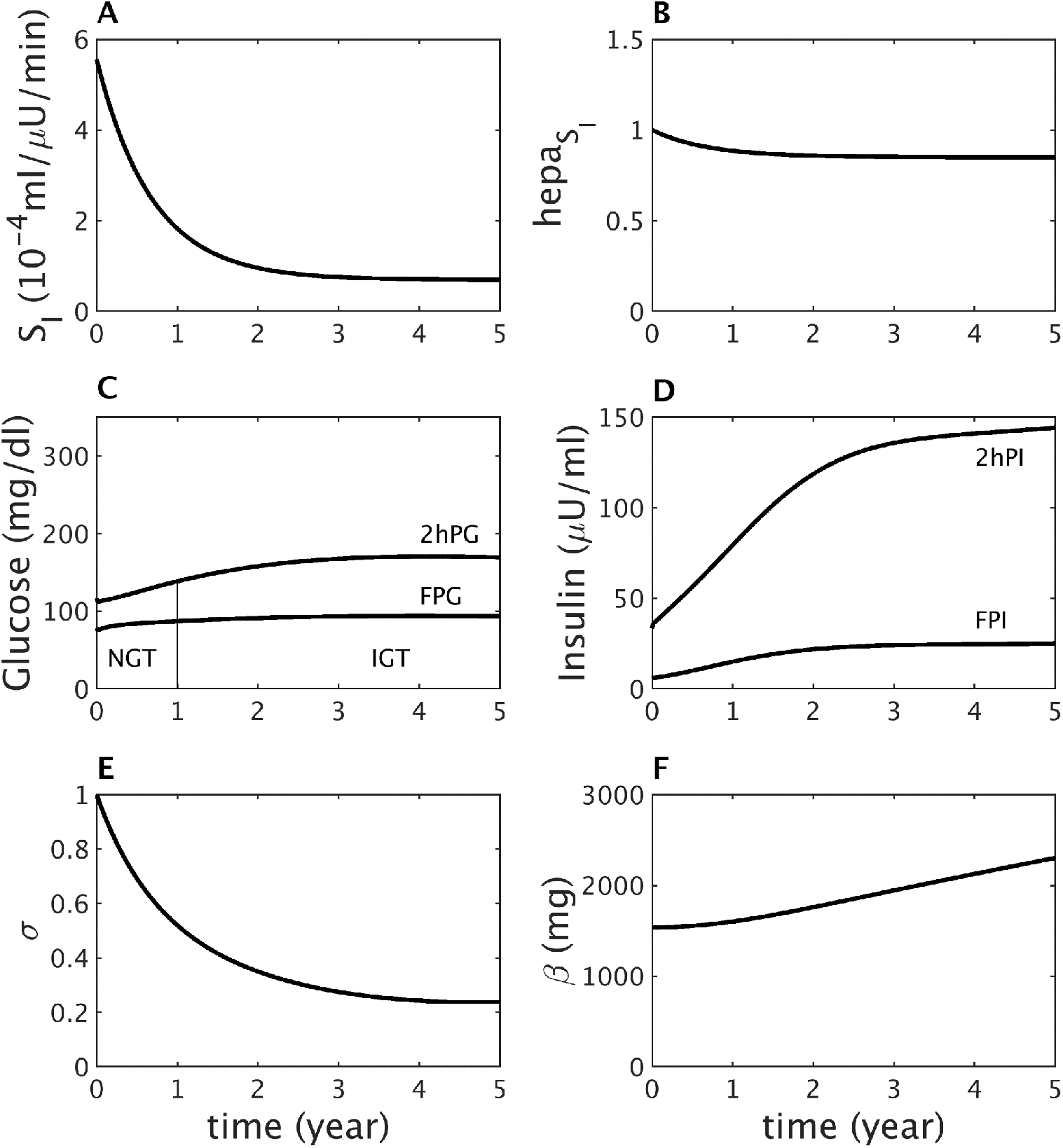
Insulin resistance (same as Fig. 1) does not lead to diabetes if beta-cell function is sufficiently responsive. The *γ*-dynamics is made stronger than in Fig. 1 by decreasing *γ*_*s*_ to 90 *mg/dl* from 100 *mg/dl* (see other parameters in Table S12A). (A) Assumed severe decline in peripheral insulin sensitivity. (B) Assumed mild decline in hepatic insulin sensitivity. (C) Simulated longitudinal changes in FPG and 2hPG during OGTTs performed at each time point. The virtual subject experiences modest rises in glucose and crosses the threshold for IGT but never crosses the thresholds for IFG, CGI or T2D. (D) Simulated longitudinal changes in fasting plasma insulin (FPI) and two-hour insulin (2hPI) during the OGTTs. Insulin concentration increases and saturates but never declines. (E) The *β*-cell function component σ first decreases, but then levels off. Increase in the *β*-cell function component represented by *γ* (not shown) helps limit the rise in glucose, which allows the *β*-cell mass *β* to increase gradually throughout (F).

Parameters that were varied to make the figures are listed in Table S12A in the Appendix, and the initial conditions for Figs. 1 – 4 are in Table S12B.

### IGT-first pathway

Figure 1 shows a longitudinal simulation of the effects of a strong decrease in peripheral insulin sensitivity *S*_*I*_ (Fig. 1A) combined with a mild decrease in hepatic insulin sensitivity, *hepa*_*SI*_ (Fig. 1B). As insulin resistance progresses, 2hPG increases rapidly while FPG increases more slowly (Fig. 1C), resulting in progression from normal glucose tolerance (NGT) to IGT, CGI and ultimately T2D (Fig. 1C).

Fasting plasma insulin (FPI) and 2-hour plasma insulin (2hPI) rise as the beta cells initially compensate partially for the insulin resistance, then fall as the beta cells fail (Fig. 1D), following the classic “Starling law” of the pancreas (27). The initial rise in secretion results from an increase in beta-cell sensitivity to glucose (the variable *γ* increases, not shown), and the decline results from a fall in the slower component of beta-cell function, σ (Fig. 1E). Beta-cell mass (*β*) also rises and falls, but the variation is limited because of the slowness of *β*, and the fall occurs only after T2D is already underway (Fig. 1F). This accords with observations that beta-cell mass is elevated in insulin-resistant pre-diabetics but reduced in long-standing diabetes (15, 66, 69).

A subtle but important point of this simulation is that insulin resistance in both the liver and peripheral tissues reaches saturation (Fig. 1A, B) well before the advent of T2D. It is rather the continuing fall in beta-cell function, σ that drives conversion to T2D. The same sequence was seen in the simulation of T2D progression in Zucker diabetic fatty rats (Fig. 6 in (37)) and in data from monkeys (10).

The fall in σ is triggered by the hyperglycemia and glucotoxicity that follows the early loss of insulin sensitivity (Fig. 1D) but would not lead to T2D if the pre-existing capacity of beta-cell function to compensate were stronger. This is illustrated by a simulation with the same degree of insulin resistance as in Fig. 1, but a milder beta-cell defect, which mimics a non-diabetic subject with insulin resistance (Fig. 2). FPG and 2hPG increase modestly in response to insulin resistance and reach a plateau in the IGT state as σ levels off (Fig. 2E) and plasma insulin plateaus (Fig. 2D). The limitation of the rise in glucose reduces the effects of glucotoxicity and buys time for beta-cell mass to increase and stabilize the IGT state (Fig. 2F).

Figures 1 and 2 together paint a picture in which not only is a combination of insulin resistance and impaired secretion necessary for T2D, but insulin resistance develops and saturates first, and T2D develops only if the beta cells fail. Some defect in beta-cell function is required even for pre-diabetes, in agreement with (28). Conversely, a pre-existing defect in insulin secretion would be silent in the absence of insulin resistance (Fig. S8), unless it were severe (Fig. S9).

### IFG-first pathway

In Fig. 3 we illustrate a contrasting case to Fig. 1, dominant hepatic insulin resistance with minor peripheral resistance (Fig. 3A, B); the same beta-cell function defect is assumed as in Fig. 1. Hepatic insulin resistance drives FPG across the threshold for IFG, while 2hPG remains below the threshold for IGT (Fig. 3C). After the initial threshold crossing, however, FPG and 2hPG continue to rise, and IFG progresses to CGI as in Fig. 1. Insulin again rises with the help of *γ* (not shown) and falls when the drop in σ becomes too great (Fig. 3E). As in the IGT-first pathway, the conversion to T2D is driven mainly by reduced beta-cell function, σ, because insulin resistance in both the liver and peripheral tissues saturates well before T2D (or even CGI) begins. Beta-cell mass again plays a minor role (Fig. 3F).

**Figure 3:**
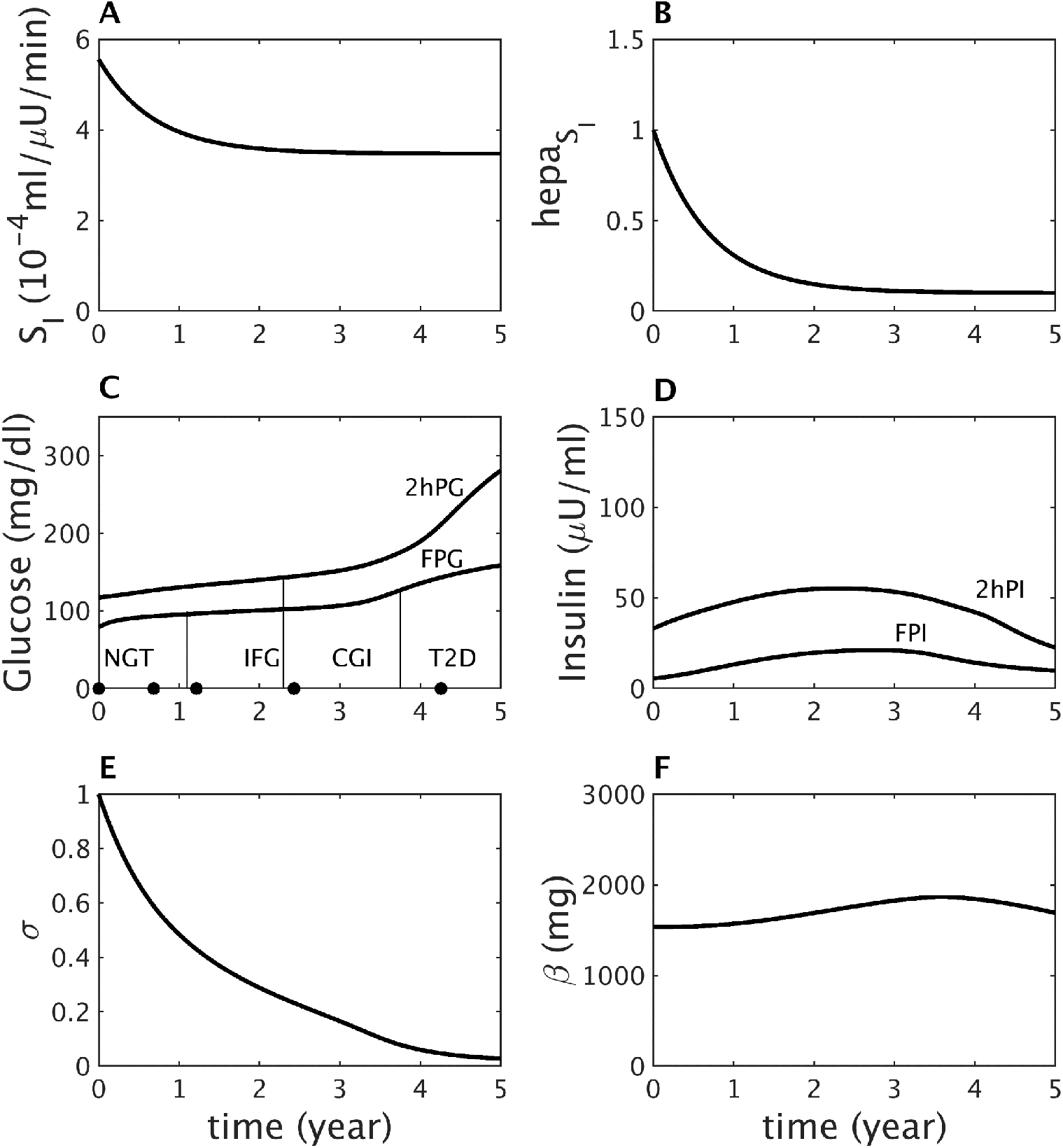
IFG-first pathway to diabetes. (A) Assumed mild decline in peripheral insulin sensitivity. (B) Assumed severe decline in hepatic insulin sensitivity component *hepa*_*SI*_. (C) Simulated longitudinal changes in FPG and 2hPG during OGTTs performed at each time point. The virtual subject experiences first high FPG (IFG), then high 2hPG (CGI), and finally crosses the FPG threshold for T2D first. Simulated longitudinal changes in fasting plasma insulin (FPI) and two-hour insulin (2hPI) during the OGTTs. Insulin concentration increases early on but decreases later. (E) The *β*-cell function component σ decreases progressively throughout. (F) The *β*-cell mass *β* increases during prediabetes then decreases after diabetes onset.

In the BLSA some subjects who had progressed from NGT to IGT went on to T2D at the next follow-up, which prompted the authors to ask whether CGI could be skipped (51). Figure 4 demonstrates that this can happen if peripheral insulin resistance is made much greater than hepatic resistance (compare Fig. 4A to Fig. 1A). Extreme loss of peripheral insulin sensitivity causes 2hPG to rise dramatically, while FPG remains in the normal range, converting NGT to IGT. 2hPG continues to deteriorate without a substantial increase in FPG, resulting in progression of IGT to T2D without passing through CGI. Eventually, FPG crosses the thresholds for IFG and T2D, but after the individual has already reached T2D based on 2hPG.

**Figure 4:**
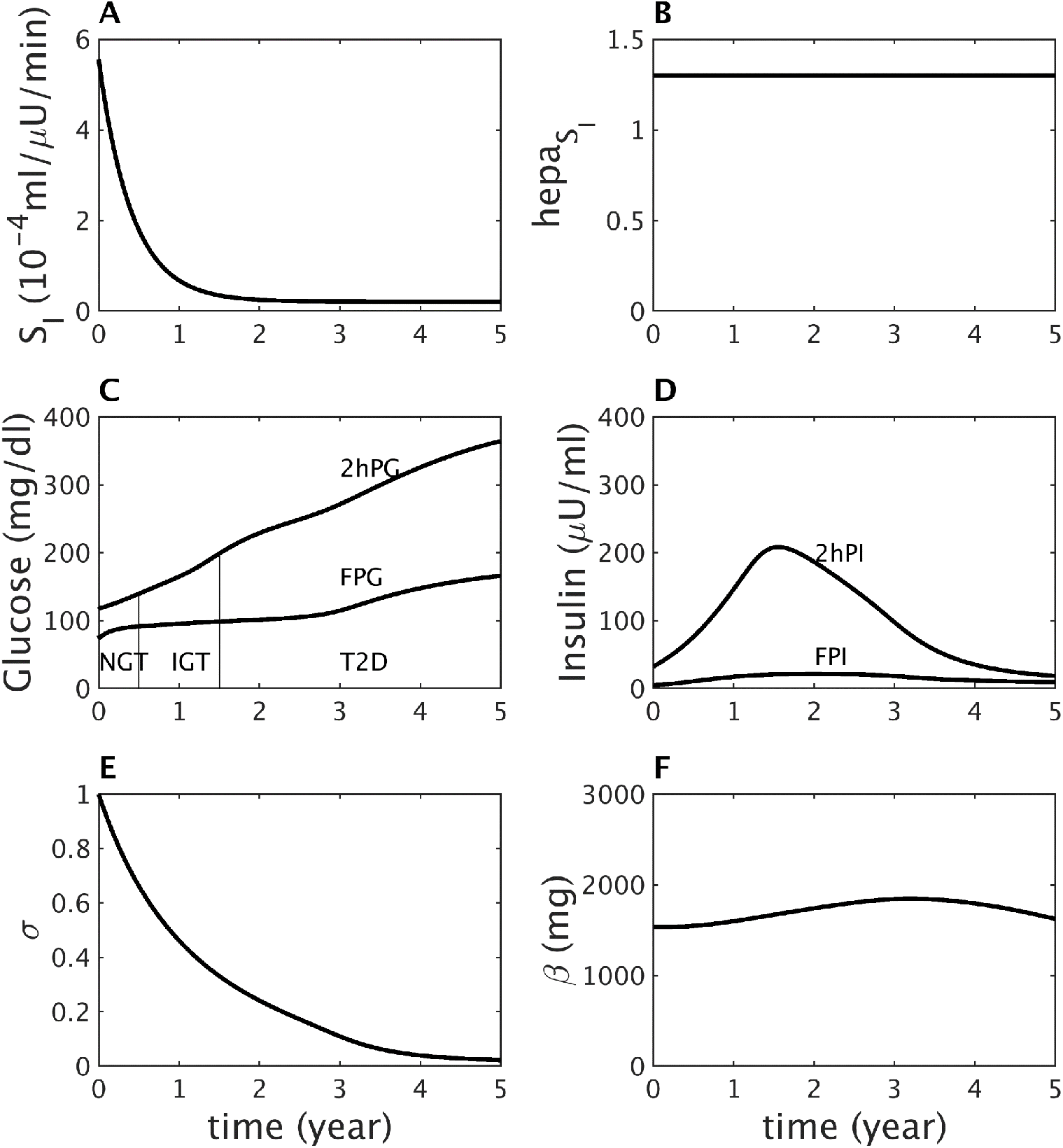
The CGI state is not obligatory. (A), (B): Extreme discrepancy between peripheral and hepatic (A – B) insulin resistance results in progression directly from IGT to T2D without passing through CGI based on OGTTs performed at each time point (C). FPG does not exceed the threshold for IFG until after T2D onset. (D) Simulated longitudinal changes in fasting plasma insulin (FPI) and two-hour insulin (2hPI) during the OGTTs. Insulin increases early on, then decreases. (E) The *β*-cell function component σ decreases more rapidly than in Figs. 1, 3. (F) The *β*-cell mass *β* increases during prediabetes, then decreases after diabetes onset.

Figures 1 and 3, respectively, consider extreme cases of peripheral insulin resistance (PIR), where *S*_*I*_ dominates, and hepatic insulin resistance (HIR), where *hepa*_*SI*_ dominates. However, most people on the path to T2D will have both. A composite view of the cases of Figs. 1 and 3 together with intermediate phenotypes is given in Fig. 5, where trajectories are plotted in the FPG – 2hPG plane. The only difference among the trajectories is the degree of HIR and PIR, varied inversely from lower right to upper left; the compensatory capacity of beta-cell function is identical for all traces. Fig. 5A shows that the trajectories diverge markedly as the defects in HIR and PIR set in but converge as hyperglycemia worsens; this happens because IFG induces IGT and IGT induces IFG. Thus, all the virtual patients end up looking the same as time goes on. The latter part of the NGT stage and the early part of the IGT stage, shown expanded in Fig. 5B, are when the underlying pathologies give rise to the most distinct behavior. This is important both for designing clinical studies and for stratifying patients for treatment. The figure suggests that the slope of the trajectory from two or more OGTTs spaced suitably far apart in time could give a good indication of the future path of the patient.

**Figure 5:**
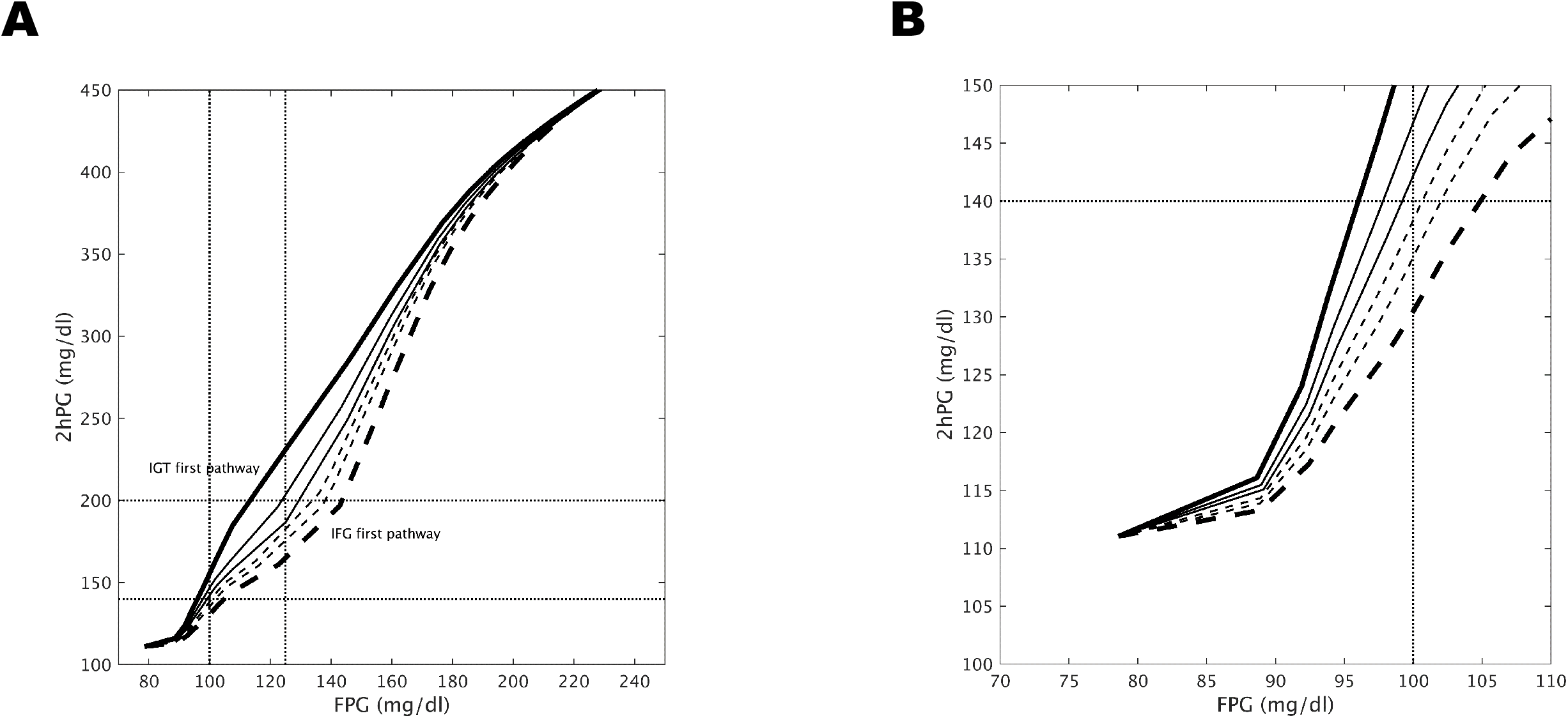
(A) Two-hour glucose (2hPG) plotted vs. fasting glucose (FPG) for varying degrees of peripheral insulin sensitivity, *S*_*I*_, and hepatic insulin sensitivity *hepa*_*SI*_, starting from the same initial values as in Figs. 1 – 4 and evolving to the target values, *tar*_*SI*_ (Eq. A13) and *tar*_*hepaSI*_ (Eq. A14) as follows: from upper left to lower right, the *tar*_*SI*_ values increase: 0.1, 0.17, 0.22, 0.33, 0.4 and 0.5; and the *tar*_*hepaSI*_ values decrease: 0.85, 0.6, 0.45, 0.35, 0.25 and 0.1. All other parameters are fixed. (B) expanded view of (A) to highlight the prediabetes region.

We next look more closely at the pathogenesis process as it would appear clinically by simulating OGTTs (Fig. 6) and IVGTTs (Fig. 7) at representative times for each stage of glucose tolerance, indicated by the black circles on the time axes in Figs. 1C and 3C.

**Figure 6:**
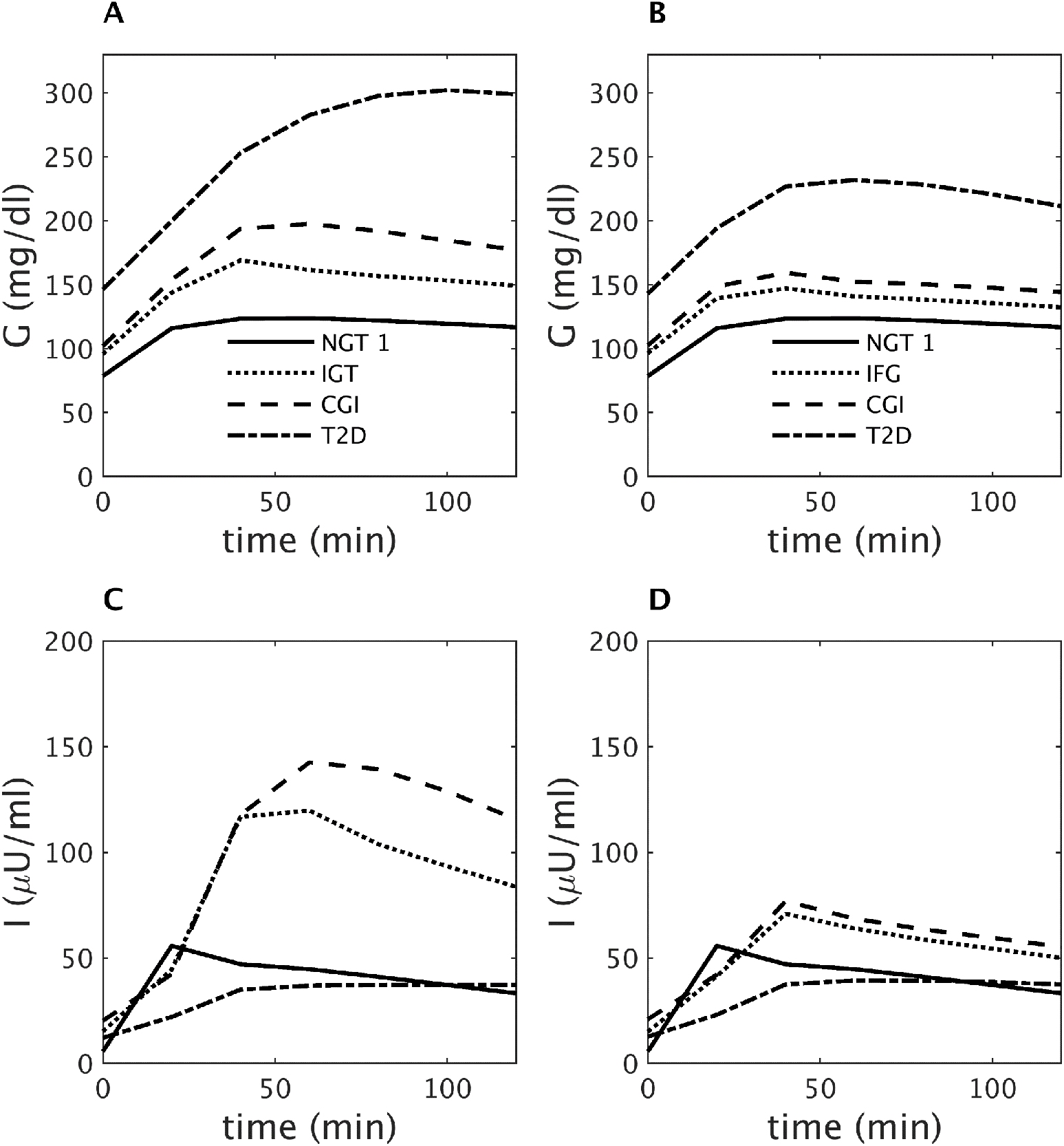
Glucose during OGTTs performed at the times indicated by the black dots in (A) Fig. 1 (IGT-first pathway) and (B) Fig. 3 (IFG-first pathway). (C, D) insulin corresponding to A and B, respectively. See text for details.

**Figure 7:**
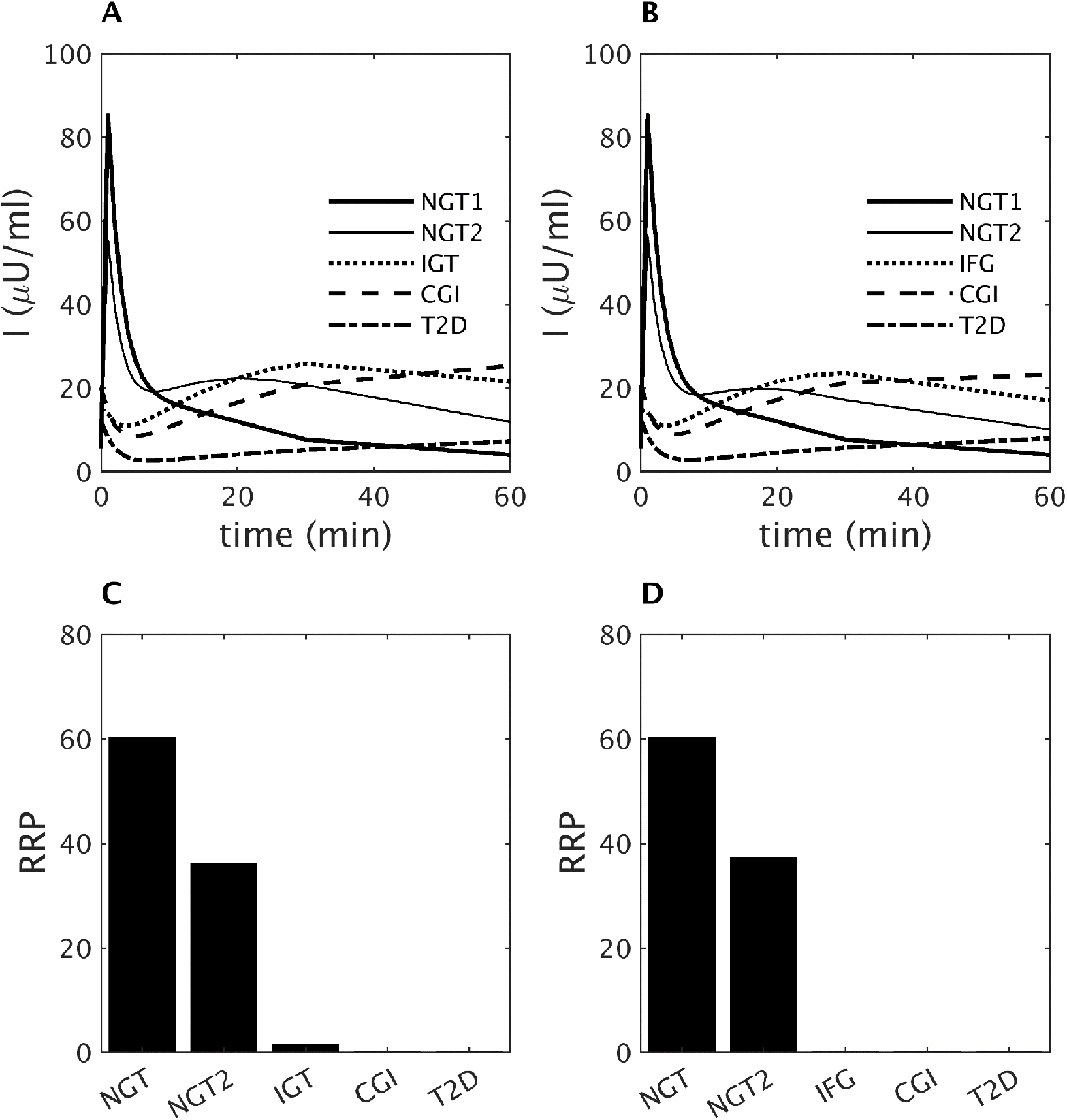
Insulin during IVGTTs performed at the time points indicated by the black dots in (A) Fig. 1 (IGT-first pathway) and (B) Fig. 3 (IFG-first pathway). (C, D): RRP size (numbers of vesicles) corresponding to A and B, respectively. Decline of AIRg parallels the reduction of RRP for each case.

### Simulation of OGTTs

Figure 6A, C shows representative glucose and insulin profiles during OGTTs at each stage of the IGT-first pathway of Fig. 1. Insulin concentrations at the IGT stage (Fig. 6C, dotted curve) are increased compared to NGT (Fig. 6C, solid curve) but are inadequate to maintain normoglycemia at two-hours because of the decreased peripheral insulin sensitivity (Fig. 1A). In contrast, the level of fasting insulin at IGT (Fig. 6C, dotted curve) is sufficient to maintain fasting glucose within the normal range because hepatic insulin resistance is relatively mild (Fig. 1B). During CGI, glucose (Fig. 6A, dashed curve) is increased at all time points compared to IGT, while the insulin level (Fig. 6C, dashed curve) is slightly diminished at the early time points (relative to glucose, however, secretion is impaired at all time points). This indicates that even progression to CGI from IGT is mainly due to impaired beta-cell function. Further and more marked decreases in insulin, due to impaired secretion at all time points during the OGTT, lead to diabetes (dot-dashed curve). Thus, during the progression towards T2D, insulin at all time points during the OGTT first increases and then decreases, in accord with the Starling Law (27).

Figure 6B, D shows glucose and insulin at each stage of the IFG-first pathway of Fig. 3. Severe hepatic insulin resistance increases glucose at all time points during the OGTT (Fig. 6B). The increase in FPG is greater than for 2hPG, so the threshold for IFG is crossed first in Fig. 3. Even though fasting insulin at the IFG stage (Fig. 6D, dotted curve) is very high, it is not enough to maintain normal FPG, because of the severe hepatic insulin resistance (Fig. 3B). However, 2hPG is maintained in the normal range because peripheral insulin resistance is mild (Fig. 3A). Since both peripheral and hepatic insulin sensitivity saturate before the onset of CGI (Fig. 3A, B), the decrease in insulin at all time points during the OGTT (Fig. 6D, dashed curve) due to falling beta-cell function (Fig. 3E) is the main contributor to the progression to CGI from IFG and then to T2D.

### Simulation of IVGTTs

Figures 1 – 6 highlight the importance of secretion defects, in the context of insulin resistance, in all the pathways to T2D, but now we break out the contributions of early vs. late secretion. First-phase secretion is widely considered a key early marker of future progress. For example, the classic paper (13) reported a cross-sectional study of IVGTTs, and showed that AIRg declines as FPG rises and is nearly gone by the time FPG reaches 115 mg/dl, well below the threshold for T2D. This supports the use of AIRg, and by implication first-phase insulin secretion, as an early marker for T2D. We now show that the model can reproduce the negative correlation between FPG and AIRg, but that rising FPG is not necessarily the sole or proximal cause of the decline in AIRg.

Figures 7A, B show simulations of the insulin responses during IVGTTs performed during the IGT-first and IFG-first pathways, respectively. AIRg is blunted and then vanishes in both pathways as FPG rises, as found in (13), but 2hPG also rises at the same time, albeit to different degrees relative to FPG in the two pathways. The decline of AIRg is more rapid than seen experimentally along both pathways (14), but the main point is that it results from the decline of RRP size (Figs. 7C, D), which is more fundamental than the level of glycemia, as the next two paragraphs explain.

RRP size is controlled by two factors, the rate of secretion, which determines the rate of vesicle efflux from the RRP and the rate of vesicle influx into the RRP from the docked pool. High FPG increases the rate of efflux in the basal state, so the RRP will already be depleted when the IVGTT commences, and AIRg will consequently be reduced. The rate of influx depends on the size of the docked pool and the rate of priming of docked vesicles. The rate of priming does not vary much in our simulations, but the size of the docked pool does, depending mainly on the rate of docking, which is proportional to one of our beta-cell function variables, σ. Although modest increases in glucose stimulate vesicle docking, larger increases cause glucose toxicity, which reduces σ and hence docking.

Because σ is slow, it responds to the average daily glucose, including the contributions of both fasting and post-prandial glucose, which differ according to pathway. Along the IGT-first pathway (Figs. 1 and 7A, C), AIRg initially declines primarily because of reduced σ (Fig. 1E), which is mainly determined by high post-prandial glucose, rather than FPG. Positive feedback in turn drives post-prandial glucose higher as σ decreases (Figs. 1C, E). Thus, the loss of AIRg and first-phase secretion is an indirect consequence of diminished second-phase secretion capacity. As the subject progresses to CGI, FPG also rises and further reduces AIRg due to pool depletion. In contrast, during the IFG-first pathway (Figs. 3 and 7B, D), the early rise of FPG reduces AIRg by depleting the RRP, and positive feedback from the later rise of 2hPG further diminishes AIRg by driving down σ.

In summary, reduced AIRg can be a marker of impairment in either first- or second-phase secretion, and, as glycemia progresses towards T2D, is likely to indicate both.

### Targeted Drug Therapy

With the previous, simpler model (37) we showed that NGT and T2D were bistable states separated by a threshold. This accounted for the well-known observation that it is easier to prevent T2D than to reverse it. The new model retains these characteristics but raises the possibility that the response to therapeutic interventions may vary depending on which pathway a patient follows to T2D. Figure 8 shows that this is indeed the case, and that knowledge of a patient’s subtype of insulin resistance can in principle lead to more effective drug therapy.

**Figure 8:**
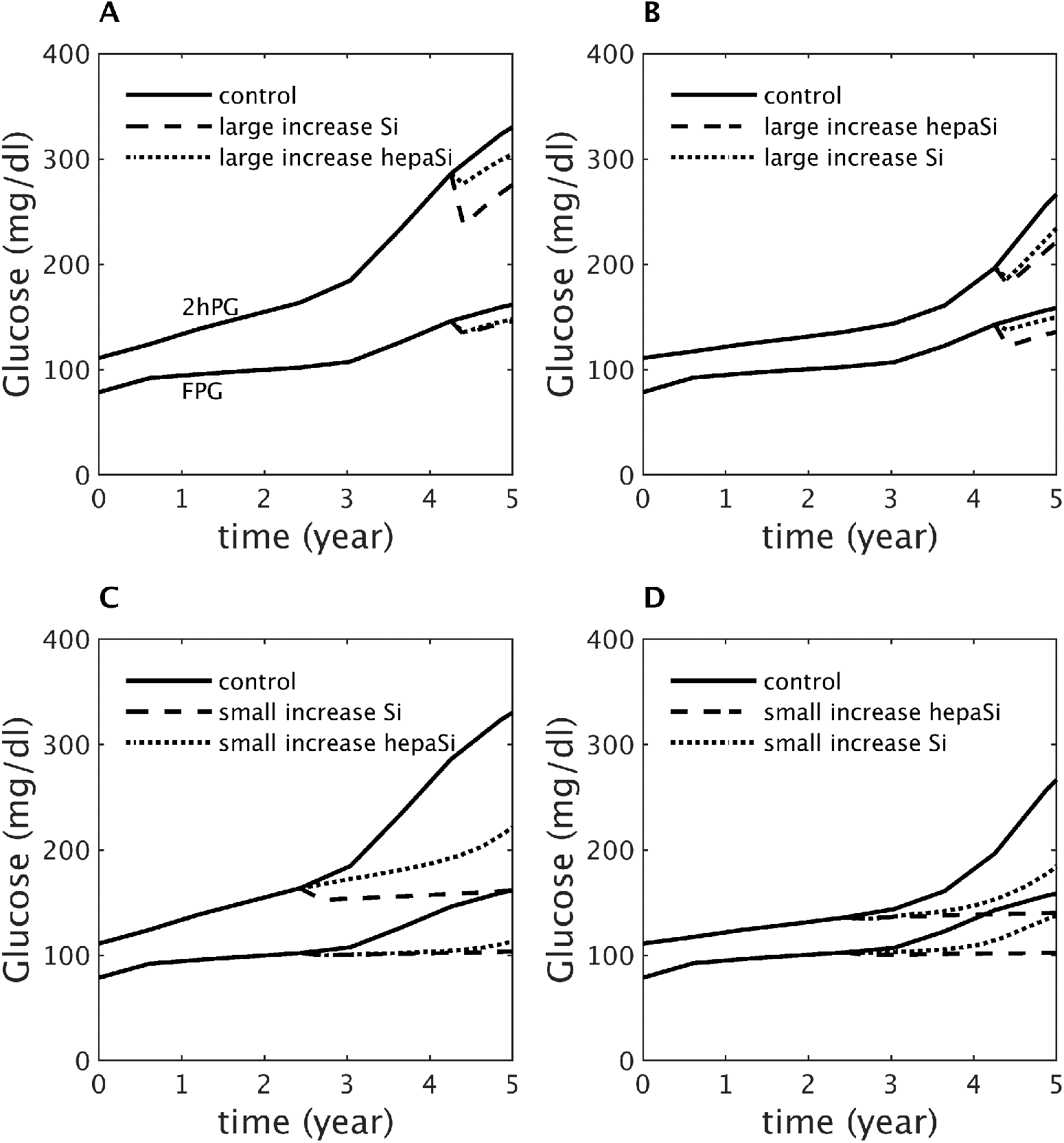
Drug therapies targeting either peripheral insulin sensitivity (modeled as a rapid increase in *S*_*I*_) or hepatic insulin sensitivity (modeled as a rapid increase in *hepa*_*SI*_; see Tables S13 – S16 for details). The drugs are applied in the early stages of T2D (A, B) or during prediabetes (C, D). (A, C) Dominant peripheral insulin resistance, leading to IGT-first pathway as in Fig. 1. (B, D) Dominant hepatic insulin resistance, leading to IFG-first pathway as in Fig. 3. The appropriately targeted drug is more effective than the inappropriately targeted drug in each case, but therapy is more effective when initiated during prediabetes.

Figures 8A, B contrast two drug therapies targeted to peripheral vs. hepatic insulin resistance in patients in the early stages of diabetes. Figure 8A shows glucose for a patient on the IGT-first pathway with dominant peripheral insulin resistance as in Fig. 1, in the absence of therapy (control, black curves) or in response to a high dose of a drug targeted to peripheral insulin resistance (dashed curves) or a drug targeted to hepatic insulin resistance (dotted). The high dose of the appropriately targeted drug only transiently improves FPG and 2hPG and ultimately fails to reverse T2D. Nonetheless, it is more effective at delaying progression than the mistargeted drug.

Figure 8B represents the complementary case, a patient on the IFG-first pathway with dominant hepatic insulin resistance, as in Fig. 3. The same treatments are applied, and, as in panel A, both drugs only transiently improve FPG and 2hPG, but the appropriately targeted drug is more effective at delaying progression.

Figures 8C, D show the same drug therapies and progression pathways as in panels A and B, but with the drugs applied before the onset of diabetes. A low dose of a drug targeted to the patient’s specific insulin resistance pathology is in both cases now able to prevent progression to diabetes, while a low dose of a mistargeted drug only delays progression. These examples suggest that the effectiveness of drug therapy depends on both early initiation of treatment and detection of the major metabolic abnormality.

The study in (24) found that it was necessary to assume that the efficacy of treatment wanes with time in order to fit the data from lifestyle and drug interventions in the DPP. Here we have shown that even if the efficacy of treatment is maintained, the intrinsic dynamics of progressive beta-cell dysfunction can cause treatment to fail.

Caution should be used in interpreting the simulated treatments in Fig. 8 in terms of currently used drugs. For example, in the DREAM study (29) it was found that rosiglitazone was effective in cases of isolated IFG (IIFG), that is IFG in the absence of IGT. Although rosiglitazone is often thought of as primarily improving peripheral insulin sensitivity, it also improves hepatic insulin sensitivity (81). In addition, IIFG does not necessarily imply pure hepatic insulin resistance. For example, a person with FPG = 115 and 2hPG = 135 would be classified as IIFG but may have significant peripheral insulin resistance. Most individuals with pre-diabetes likely have a mixture of peripheral and hepatic insulin resistance.

### Hepatic Insulin Clearance

Another way to generate a compensatory increase in insulin is to reduce clearance. There is considerable evidence that insulin clearance is positively correlated with peripheral insulin sensitivity (3, 63), but the direction of causation is not clear. If clearance is reduced secondary to reduced overall insulin sensitivity, then the increased insulin would contribute to the compensatory response, along with increased secretion. However, it has been suggested that the increased insulin resulting from reduced clearance may also contribute to insulin resistance (61). In Fig. 9, we explore the consequences of reduced clearance, assuming that the increased insulin reduces *S*_*I*_ is reduced by varying degrees as *I* is increased, as described in Methods. We assume for simplicity that clearance itself is constant throughout each simulation rather than adapting to glucose, insulin or insulin sensitivity. The control case, shown in black, is equivalent to Fig. 1, an example of an IGT-first pathway to T2D. If clearance is reduced, and a modest degree of additional insulin resistance is assumed (green curve in panel F), such that the *S*_*I*_ is only slightly lower than the control case (panel A), then the progression to T2D is delayed (compare black and green curves in panel C). (The smallness of the difference in *S*_*I*_ is due to the fact that glucose is elevated to levels where the induced effect is triggered only for a brief period of each day.) Note that σ is slightly larger (green vs. black, panel E), indicating that reducing clearance partially spared beta-cell function.

**Figure 9:**
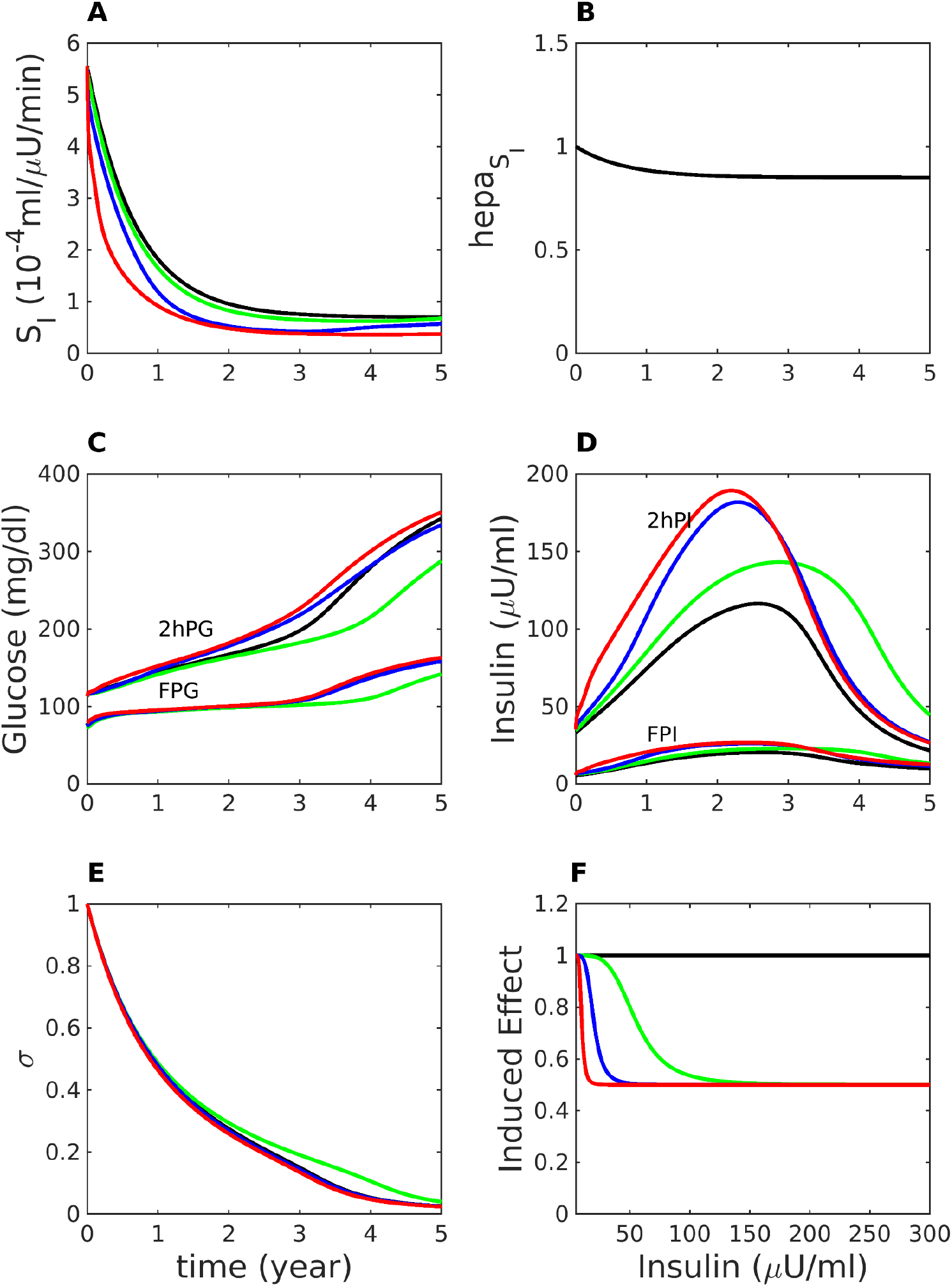
Effects of insulin clearance and insulin-induced insulin resistance. The case of Fig. 1 (IGT first) is re-simulated with varying rates of insulin clearance and induced insulin resistance, defined by Eqs. 13 – 16. Panel A: *S*_*I*_ = *S*_*I*,0_*E*, where *S*_*I*,0_ is calculated as in Fig. 1 and *E* is the induced effect plotted in Panel F. Legend: *black* (control), same as Fig. 1, but with insulin clearance *k* = 0.4861 min^−1^, essentially no induced effect (*α*_*E*_ = 300 μU/ml); *green*: reduced clearance (*k* = 0.37499 min^−1^), modest induced effect (*α*_*E*_ = 50 μU/ml); *blue*: *k* = 0.37499 min^−1^, stronger induced effect (*α*_*E*_ = 15 μU/ml); *red*: *k* = 0.37499 min^−1^, very strong induced effect (*α*_*E*_ = 5 μU/ml). Other parameters are listed in Table S17.

If the effect of insulin to induce resistance is made larger (blue curves), the beneficial effect of higher *I* due to reduced clearance and the harmful effect of reduced insulin sensitivity approximately cancel out. Finally, if the induced resistance effect is made extreme, such that even a slight increase in *I* reduces *S*_*I*_ in half (red curves), the net effect is somewhat accelerated progression. We conclude that induced insulin resistance makes at best a modest contribution to the progression to T2D. If an even more extreme degree of induced resistance is assumed, such that even a slight increase in *I* reduces *S*_*I*_ to near 0, then it is possible for frictional resistance to be the main driver of T2D, but we consider such a scenario implausible.

### Non-Insulin-Mediated Glucose Uptake

We have focused up to this point on the contributions of hepatic and peripheral insulin resistance in driving the IFG-first and IGT-first pathways. However, non-insulin-mediated glucose uptake, also known as glucose effectiveness, is comparable in magnitude to insulin-dependent uptake (9) and has been found to be impaired in T2D (30). It is thus a potential target of drug therapy. We model this by varying the model parameter *E*_*G0*_ (similar to *S*_*g*_ in Minimal Model analysis) in Eq. 1. In Fig. S10 we demonstrate that cutting glucose effectiveness in half can result in T2D in someone otherwise not at risk, such as the case shown in Fig. 2. Conversely, in Fig. S11 we show that doubling glucose effectiveness can prevent T2D in someone otherwise predisposed, such as the case shown in Fig. 1.

## Discussion

In recognition of the fundamental character of type 2 diabetes (T2D) as a progressive disease that develops over many years, we have established a longitudinal model for its pathogenesis. We follow in the footsteps of other longitudinal models (23, 24, 75) but offer new clinical applications and insights.

We apply the model to analyze the diverse presentation of hyperglycemia, which may manifest first in fasting glucose (IFG-first pathway) or two-hour glucose during an OGTT (IGT-first pathway). To carry out this program, we modified the representation for beta-cell response to insulin resistance in the first version of the model (37), enhancing it to differentiate between hepatic and peripheral insulin resistance. The simulations show that heterogeneity in the degree of the two forms of insulin resistance can account for a wide variety of observed patterns, supporting the idea of T2D as a unitary disease with quantitative variants. We have focused on extreme cases to highlight the differences (e.g. Fig. 1 vs. Fig. 3), but the family of trajectories in the FPG-2hPG plane (Fig. 5) shows that these lie on a continuum. Figure 5 also highlights that differences in insulin resistance phenotype are most evident in the late NGT and early pre-diabetes stages, which are thus most amenable to differential phenotyping and therapeutic stratification.

We have incorporated a description of insulin granule dynamics sufficient to account for both first- and second-phase insulin secretion. This made it possible to simulate OGTT and IVGTT time courses and show how they are transformed systematically during progression along the two canonical pathways to diabetes (Figs. 6, 7). We also showed that sufficiently strong beta-cell function can prevent T2D even when insulin resistance is severe, allowing individuals to maintain a permanent state of IGT (Fig. 2) or even revert from IGT to NGT (not shown). Conversely, sufficient insulin sensitivity can prevent T2D even when beta-cell function is somewhat impaired (37). As discussed below under *Secretion defects*, a fuller treatment of the differences in the balance of insulin secretion and insulin action defects is needed to account fully for the diverse patterns of T2D progression.

We summarize below the specific lessons learned and questions answered by this study and give a preview of the clinical applications we anticipate for the model.

### Questions raised in the BLSA and other studies

The BLSA study (51) asked whether subjects who enter the IGT state necessarily pass through CGI on the way to T2D. The model suggests (Fig. 4) that this is not the case, but skipping CGI happens only if the peripheral insulin resistance is markedly greater than hepatic insulin resistance, which may be relevant for adolescents, who experience extreme obesity and insulin resistance. An intermediate possibility predicted by the model is a short, but not absent, interval of CGI that could escape detection if the follow-up interval is too long.

A parallel question is whether individuals can go directly from IFG to T2D without passing through CGI. This is harder than going directly from IGT to T2D because 2hPG is much more labile than FPG; it is difficult to get an increase in FPG sufficient to cross the threshold for T2D (125 mg/dl) without at the same time having 2hPG cross the threshold for IGT (140 mg/dl). Indeed, we have not been able to simulate this scenario just by choosing an appropriate mixture of hepatic and peripheral insulin sensitivity using the other parameters as in Figs. 1 – 4, but model simulations (not shown) predict that it can happen if a more severe beta-cell defect (in *γ*_∞_ Eq. A15) is assumed.

The BLSA study also asked whether individuals can pass directly from NGT to T2D without passing through any pre-diabetic state. Because glucose is in quasi-steady state with the much slower variables representing beta-cell mass, beta-cell function and insulin sensitivity, this is not possible in the model unless one of those slow variables undergoes a catastrophic, virtually discontinuous, change. Pancreatectomy would be an example of this, but even type 1 diabetes, triggered by a rapid fall in beta-cell mass, has a distinct prediabetes phase. The cases observed in the BLSA in which subjects were NGT at baseline and T2D at first follow-up most likely reflected rapid progression or a long gap between visits. Similarly, the model simulations indicate that it is unlikely for individuals to go from NGT to CGI without passing through IFG or IGT.

The BLSA reported that IFG is generally followed by IGT and IGT is generally followed by IFG, and the model suggests that each state *induces* the other. This happens because the initial rise in glucose during IFG impairs beta-cell function, which causes 2hPG to rise, and vice versa. Another study raised the question of whether CGI is a progressed state of IFG (57); the model simulations together with the BLSA data show that CGI may instead be a progressed state of IGT, to which IFG has been added.

One can also ask whether crossing the threshold for FPG or 2hPG of pre-diabetes predicts whether T2D will be reached by crossing the corresponding threshold. In Fig. 1, IGT is followed by T2D diagnosed through 2hPG, and further simulations with the model (Fig. 5) suggest that this is typical. In Fig. 3, IFG is followed by T2D diagnosed through FPG, but further simulations (not shown) indicate that this may or may not be the case, depending on the degree of discrepancy between HIR and PIR and the strength of *γ* to control FPG.

Both the IFG- and IGT-first pathways exhibit elevated fasting insulin. This may account for the observation that elevated fasting insulin was a better predictor of future diabetes in a prospective study than fasting glucose, which is elevated early on only in the IFG-first pathway (22); it may not be necessary to hypothesize a major causative role for high fasting insulin itself. Indeed, in the IGT-first pathway, which is more common, fasting glucose may be suppressed by the compensatory increase in beta-cell function (our variable σ) induced by high post-load glucose (compare Figs. 1 and 4 to Fig. 3).

The suppression of fasting glucose in the context of a predominance of peripheral insulin resistance has particular importance for pre-diabetes screening in populations that are prone to IGT but not IFG. This applies notably to people of African descent, for whom fasting glucose has markedly reduced sensitivity for detecting pre-diabetes and diabetes (77). The problem is exacerbated for Africans living in Africa, where measuring 2hPG with OGTTs is prohibitively expensive. The model suggests that in this and similar cases, lowering the threshold for diagnosing pre-diabetes based on fasting glucose could be a cost-effective strategy. More generally, the model points to the need for population- and patient-specific thresholds for diagnosis, which may contribute to resolving current debate on whether prediabetes is a useful diagnosis (62).

### Insulin action defects

We have varied peripheral and hepatic insulin resistance independently to study their contributions to T2D progression, but in reality, they are related. Statistically, they are correlated with a coefficient of about 0.7 (1). This is expected for several reasons. For one, they share major components of the insulin signaling pathway. Also, there is evidence that an important determinant of hepatic insulin resistance is excess supply of free fatty acid (FFA) substrate from adipose tissue (58). Thus, insulin resistance in adipose cells would increase lipolysis and FFA flux to the liver, which would drive increased gluconeogenesis (59). On the other hand, the liver has unique roles in glucose and lipid production not shared with muscle, which may account for the fact that the correlation is imperfect.

To address hepatic insulin resistance above and beyond the component correlated with peripheral insulin resistance, we have independently varied the affinity of HGP for insulin (parameter *hepa*_*SI*_ in Eq. 9), the effect of which is shown in Fig. S4. We have obtained similar results (not shown) by varying the maximal rate of HGP (parameter *hepa*_*max*_).

Although we have primarily treated insulin resistance as an external influence on the model, representing genetic and obesity, we have considered the much discussed hypothesis (65, 71, 74) that hyperinsulinemia can contribute to insulin resistance in addition to being a compensatory response to insulin resistance (Fig. 9). Any effect of increased insulin to induce insulin resistance has to be weighed against the effect to enhance glucose uptake, and the outcome would depend on the net effect on the term *S*_*I*_*I* in Eq. 1. We find that incorporating induced insulin resistance can either advance or delay the rate of progression to T2D modestly depending on the strength of the two effects. We have not found support for the hypothesis that it is the sole or primary initiating factor: to trigger T2D in the absence of intrinsic, insulin-independent insulin resistance, we had to assume that the insulin-induced resistance is implausibly strong. Similar results would apply if hyperinsulinemia resulted from increased secretion rather than reduced clearance, but one would also have to consider whether in this case the increased beta-cell workload might have an additional deleterious effect.

We are aware of only one other modeling study (35) that has addressed the role of hyperinsulinemia as a driver of diabetes pathogenesis. More specifically, that thoughtful analysis considered a variant of the model of Topp et al (75) in which *hypersecretion* would reduce *S*_*I*_. Under that assumption, and an additional assumption that high insulin would impair beta-cell mass, hypersecretion was found to be able to drive progression to T2D. However, increased insulin itself that has been shown to induce insulin resistance (71), so we prefer to have insulin act directly on *S*_*I*_. As we have said above, the effect of hyperinsulinemia governed by that mechanism is modest.

The effects of induced insulin resistance are limited fundamentally because insulin both promotes glucose uptake and contributes to resistance to that uptake. This is analogous to friction in mechanics. The higher the insulin concentration, the more resistance is generated through increased negative feedback in the insulin signaling cascade. Similarly, the faster a car goes, the more wind and rolling resistance it generates. This resistance is not the motive force driving the car forward but a by-product of forward motion that limits its speed. However, resistance is generally not strong enough to prevent acceleration, it just requires more gas to achieve the desired speed. Similarly, the insulin resistance induced by insulin can in most cases be overcome by secreting more insulin.

### Insulin secretion defects

The simulations of progression to T2D (Figs. 1, 3, and 4) require some degree of secretion defect in addition to the various combinations of insulin resistance. We assumed a defect only in the triggering pathway (*γ*), but we have obtained similar results by assuming defects in the amplifying pathway (σ). The assumed triggering defect represents a right shift in the glucose dose response curve, which correspond to a mild gain of function (GOF) mutation of the K(ATP) channels, a prominent GWAS hit (55).

Assuming a more significant GOF mutation defect in the model leads to T2D with a lesser degree of insulin resistance (Fig. S8). This is in line with the finding that K(ATP) defects are generally mild (for example, the E23K mutation in the Kir6.2 component of K(ATP), KCNJ11, has an odds ratio of only 1.18 (33)), but they are fairly common and therefore may make significant contributions on the population level in combination with insulin action defects.

If the K(ATP) defect is sufficiently severe, no insulin resistance is needed at all (Fig. S9). This is the case in neonatal diabetes mellitus (NDM) (4, 47), which is of particular interest because it can be treated effectively with sulfonylureas. This differs from typical T2D in adults, possibly because the drug brings the genetically insufficiently active beta cells to a normal level of activity instead of overdriving them.

The examples of Figs. 2, 1, S8 and S9, in that order, lie on a continuum of progressively greater K(ATP) channel function and lesser insulin resistance. As we showed previously in Fig. 5 of Ref. #6, the greater the degree of beta-cell dysfunction, the lesser the degree of insulin resistance required for T2D. In addition to the degree of insulin resistance, the rate of decline of insulin sensitivity is important. Fig. S12 shows cases in which a slow decline is compensated whereas a fast decline is not, as shown previously in (75). We emphasize that the requirement for insulin resistance for typical T2D progression is not in contradiction with genome wide studies showing a predominance of beta-cell genes: most people who are overweight or obese become insulin resistant, but only those with beta-cell defects become diabetic. Lean individuals who are not insulin resistant can also develop diabetes, but only if they have severe and rare secretion defects.

Although we have focused on the role of insulin resistance in promoting IFG, secretion defects can also lead to IFG with little contribution of insulin resistance. Fig. S13 shows a case with a mild GOF mutation in K(ATP) channels, and Fig. S14 shows a case with a small readily releasable pool due to reduced vesicle priming rate. With more extreme secretion defects, no insulin resistance is needed at all (not shown). These patterns may be of particular relevance to Asian and possibly African populations where secretion defects play a bigger role than insulin resistance in diabetes compared to those of European descent.

Another secretory defect that may contribute to diabetes risk is impaired incretin signaling. Figure S15 illustrates that such an impairment would accelerate progression in a person predisposed to T2D.

The model shows that defects in first- and second-phase secretion are not independent. Impairment in one leads to impairment in the other because of the harmful effects of elevated glucose. The model accounts for classic data (13) showing that the acute insulin response to glucose (AIRg), a surrogate for first phase secretion, declines with even modest increases in FPG during the NGT and pre-diabetic stages. The model shows, however, that second-phase insulin secretion declines in parallel, belying a privileged role for first-phase secretion, as also shown in another modeling study (36). In our model, reduced AIRg results mainly from reduced size of the RRP, which can be caused by impairment in either first-phase or second-phase secretion or both. More generally, any unanswered rise in either fasting or post-prandial glucose impairs both first and second phase secretion, and any defect in either first or second phase secretion raises both fasting and post-prandial glucose. This creates a vicious cycle that drives down both first- and second-phase secretion regardless of which defect is primary. Empirical studies that do not select for early isolated IFG or IGT cases do not show a clear prominence of first phase secretion loss (31), in agreement with the model.

### Non-insulin mediated glucose uptake

We have also considered briefly glucose effectiveness, the ability of glucose to promote its own uptake independent of insulin (Figs. S10, S11). For simplicity, we have assumed the differences in *E*_*G0*_ to be pre-existing and constant in time. However, there is evidence that it is a regulated process; it has been reported to be enhanced by FGF19 acting in the brain and may be related to the glucose intolerance resulting from leptin deficiency in ob/ob mice (52). It may also account for the rescue by leptin of glucose tolerance in insulin-deficient rodents treated with streptozotocin (32). Further progress in modeling this process would benefit from identifying the molecular mechanism(s) underlying it, which are currently not well understood.

### Limitations of the study

The model as presented here is oversimplified in that it only considers glucotoxicity and neglects other factors that are likely to play a role, such as lipotoxicity (48). We are thus unable at present to address the possible role of hyperinsulinemia to promote hepatic fat accumulation (76). If prolonged, this could lead to liver damage and thus to a worsening of insulin resistance that could outlast the induced resistance we have modeled here (Eqs. 9 – 11), which would resolve if normal insulin levels were restored.

In addition, we believe that prolonged high secretion rate is probably harmful beyond the negative effect incorporated in the equation for σ, possibly because of ER stress and/or calcium toxicity (53). Those factors were not included here because they were not needed to account for the pathways considered, but they may be necessary to explain other pathways to T2D and will be addressed elsewhere.

To capture the full range of patterns, it is necessary to consider as well pre-existing variation in beta-cell function, not the moment-to-moment beta-cell function, which evolves in response to hyperglycemia, but the innate, genetic capacity of beta-cell function to adapt to hyperglycemia. For example, changing the parameters defining σ_∞_ (Eqs. 6, A16) has marked effects on the speed of progression in IGT; we neglected this because it doesn’t change the likelihood of entering the IGT state much.

We have modeled the suppression of hepatic glucose production as a direct effect of insulin on the liver. However, much if not all of the acute effect of insulin is indirect, mediated by suppression of lipolysis in adipose tissue, which reduces the supply of free fatty acid (FFA) to the liver (58). We consider this an acceptable approximation for our purposes, as post-prandial suppression of lipolysis is roughly a mirror image of the post-prandial rise in insulin. In future, if we want to account for FFA dynamics or adipose-tissue insulin resistance, the model would have to be augmented.

We have modeled insulin-dependent glucose uptake as linear in insulin, but it is likely to be non-linear (sigmoidal) (12) and has been suggested to exhibit hysteresis (78). We have not found these features to be necessary to explain the data under consideration, but they can be easily added in the future if the need arises.

We have neglected the role of excess glucagon secretion, which results from intrinsic defects in alpha cells or from loss of paracrine regulation of alpha cells due to a relative lack of insulin (21, 56). This has long been recognized to exacerbate post-prandial hyperglycemia in diabetes and pre-diabetes and is a target of GLP-1-based insulin secretagogues. Without glucagon in the model, we are not able to simulate fully the efficacy of that class of drugs, only their effect on insulin secretion. Hyperglucagonemia may also contribute to hepatic insulin resistance, as proposed in recent reviews (26, 80) suggesting that glucagon is part of a liver-alpha cell axis that is primarily devoted to regulating amino acid metabolism. A primary form of glucagon resistance at the liver in this cycle could lead to hyperaminoacidemia and hyperglucagonemia. The latter would contribute to excess hepatic glucose production and IFG in addition to or in place of hepatic insulin resistance. This makes glucagon a tempting therapeutic target. Hyperaminoacidemia and alpha-cell hyperplasia have thwarted many efforts, but some work in this direction continues (72).

Other recent developments have shown that glucagon amplifies insulin secretion and thus under normal conditions has a net hypoglycemic effect post-prandially (16, 82). Glucagon is thus now seen to be homeostatic in both low and high glucose under normal conditions but can become anti-homeostatic in diabetes or pre-diabetes. The emerging story is complex, and inclusion in the model is deferred to future studies.

We have included only a very simple representation of insulin clearance, assuming a first-order dependence on insulin concentration. We have not distinguished portal from peripheral insulin (63) or considered possible regulation by glucose (60) or free fatty acids (7).

### Future directions

This paper has demonstrated that if the insulin resistance phenotype of an individual is known, their future trajectory of hyperglycemia can be predicted (Figs. 1 – 4) and drug choice can potentially be optimized for the patient’s insulin resistance phenotype (Fig. 8). Another longitudinal model has shown similarly the results of targeting insulin resistance vs. beta-cell replication (38). Our model can provide similar predictions if the relative impairments in insulin secretion and insulin resistance are known. For example, plotting 1hPG vs. 2hPG obtained by simulating OGTTs results in a family of trajectories similar to the ones for FPG vs. 2hPG shown in Fig. 5. The differences in trajectories due to the contributions of insulin secretion and insulin action can be read off from the difference between 1hPG and 2hPG and are similarly most pronounced during prediabetes (Ha et al, ADA poster 1490-P, June, 2019). This prediction may provide deeper insight into the diagnostic information that can be extracted from different time points during the OGTT, an issue that is currently receiving much attention (19, 40).

Future versions of the model should include roles for glucagon, as described above. In addition, a more complex model for insulin signaling would permit the investigation of the effects of particular defects in those pathways as well as possible mechanisms for the insulin resistance induced by insulin, which was treated phenomenologically here.

In addition, to be a useful tool for patient stratification and treatment planning, one needs to ascertain the patient’s phenotype. We plan to investigate whether the model can be used, with suitable modifications, to solve the inverse problem of inferring the individual’s parameters of insulin resistance and beta-cell function from the observed behavior.

## Acknowledgments

The work was supported by the Intramural Research Program of the National Institutes of Health, NIDDK. We thank Stephanie Chung, Richard Bertram, Over Cabrera, and Cecilia Diniz Behn for helpful suggestions on the manuscript.

## Notes

http://lbm.niddk.nih.gov/sherman

### Competing Interest Statement

The authors have declared no competing interest.

### Summary of Updates

Additional figures have been included comparing the model outputs to published experimental data and additional commentary on features omitted from the model have been made in the Discussion.

